# The APL1 immune factor is encoded by a single ancestral gene in most *Anopheles* species and expanded to three paralogs with distinct function in the *Anopheles gambiae* complex

**DOI:** 10.1101/785089

**Authors:** Christian Mitri, Emmanuel Bischoff, Karin Eiglmeier, Inge Holm, Constentin Dieme, Emma Brito-Fravallo, Abbasali Raz, Sedigheh Zakeri, Mahdokht I. K. Nejad, Navid D. Djadid, Kenneth D. Vernick, Michelle M. Riehle

## Abstract

**Background:** The recent reference genome assembly and annotation of the Asian malaria vector *Anopheles stephensi* revealed only one gene encoding the leucine-rich repeat immune factor APL1, while in *Anopheles gambiae* and sibling *Anopheles coluzzii*, APL1 factors are encoded by a family of three paralogs. The phylogeny and biological function of the unique APL1 gene in *A. stephensi* have not yet been specifically examined.

**Methods:** The APL1 locus was manually sequenced to confirm the computationally predicted single APL1 gene in *A. stephensi*, and APL1 evolution within *Anopheles* was explored by phylogenomic analysis. The single or paralogous APL1 genes were silenced in *A. stephensi* and *A. coluzzii*, respectively, followed by mosquito survival analysis, experimental infection with *Plasmodium*, and expression analysis.

**Results:** APL1 is present as a single ancestral gene in most *Anopheles* including *A. stephensi*, but has expanded to three paralogs in an African lineage that includes only the Gambiae species complex and *Anopheles christyi*. Silencing of the unique APL1 copy in *A. stephensi* results in significant mosquito mortality. Elevated mortality of APL1-depleted *A. stephensi* is rescued by antibiotic treatment, suggesting that bacteria are the cause of mortality, and that the unique APL1 gene is essential for host survival. Successful *Plasmodium* development in *A. stephensi* depends upon APL1 activity for protection from high host mortality, probably caused by exposure to enteric bacteria when parasites cross the midgut epithelial barrier. In contrast, silencing of all three APL1 paralogs in *A. coluzzii* does not result in elevated mortality, either with or without *Plasmodium* infection. Expression of the single APL1 gene is regulated by both the Imd and Toll immune pathways, while control by the two pathways is subdivided to different paralogs in the expanded APL1 locus.

**Conclusions:** APL1 underwent neofunctionalization with both loss and gain of functions concomitant with expansion from a single ancestral gene to three paralogs in one lineage of African *Anopheles*. The evolution of an expanded APL1 gene family could be a factor contributing to the exceptional levels of malaria transmission mediated by human-feeding members of the Gambiae complex in Africa.

## BACKGROUND

Malaria remains a serious global public health concern. Human malaria is transmitted by *Anopheles* mosquitoes, and among >450 extant *Anopheles* species, approximately 40 are considered dominant malaria vector species (DVS) (1). About 90% of global *Plasmodium* falciparum transmission occurs in Africa, where the most important DVS on earth are members of the *Anopheles gambiae* species complex (hereafter, Gambiae complex), including the widespread *Anopheles coluzzii*. An important Asian DVS is *Anopheles stephensi*, which has recently been recognized as an invasive vector species, expanding disease transmission along with its geographic range (2, 3).

The heterogeneity among *Anopheles* species for malaria vectorial capacity can have multiple causes. The first is host-feeding behavior, because animal-feeding species do not have the opportunity to acquire and transmit a human pathogen, and consequently human-biting preference is the most fundamental prerequisite of malaria vectorial capacity (4, 5). *Anopheles* that are designated non-vectors are not necessarily refractory to *Plasmodium*, and when fed experimentally on parasitemic blood, behavioral non-vectors can be physiologically competent for *Plasmodium* infection (6, 7).

Among human-feeding DVS, there is apparent variation in vectorial capacity, suggested by large geographic differences in human malaria infection prevalence, with about 90% of global prevalence located in Africa (8). Some of this global geographic variation could be caused by ecology, if some niches, for example in humid sub-Saharan Africa, are particularly favorable to mosquito abundance and longevity, promoting malaria transmission (9–11). Finally, vector genetic differences can also underlie physiological difference in vector competence for *P. falciparum* in nature (12–14), but the mechanisms underlying *Anopheles* susceptibility to human malaria in nature are not understood. Several tens of *Anopheles* genes are known from laboratory studies to control malaria infection of the vector, but involvement of these genes in modulating natural transmission has not been confirmed by genetic association in the natural vector population.

The best described mechanism of mosquito immunity in laboratory studies is a ternary immune complex in the Gambiae complex, comprised of the leucine-rich repeat (LRR) proteins APL1 and LRIM1, and the complement-like factor TEP1 (15–17). APL1 is present in the Gambiae complex as a family of three paralogs, APL1A, APL1B and APL1C (16). The paralogs display distinct spectra of protection for different pathogen classes (18–20). APL1A activity inhibits development of the human parasite *P. falciparum*, while APL1C activity inhibits rodent malaria species (16), and APL1B modulates protection against both *P. falciparum* and the rodent parasites (19).

The recent reference genome assembly and annotation of the Asian malaria vector *A. stephensi* revealed only one APL1 gene rather than three paralogs as in the Gambiae complex (21). Here, we experimentally validate the computationally predicted single APL1 gene in *A. stephensi*. Phylogenomic analysis indicates that a single copy of APL1 represents the ancestral anopheline state, while the expansion to three APL1 paralogs is derived, and among DVS is found only in the African lineage that includes the Gambiae complex. *A. stephensi* APL1 was previously tested for effect on *P. falciparum* (22) and response to kinase signaling (23), but the biological function of the unique APL1 gene has not yet been specifically examined, nor compared to the function of the expanded APL1 locus. We find that the single-copy ancestral APL1 gene and the expanded APL1 locus display distinct functional properties, including for host survival, protection against *Plasmodium* infection, and immune signaling. The expanded APL1 locus is found in the most efficient DVS in the world, the Gambiae complex, which poses the question whether the apparent correlation of APL1 copy number with efficient malaria transmission is accidental or causal.

## METHODS

### Phylogenetic analysis of *Anopheles* APL1 gene copy number

We confirmed by manual sequencing the existence of a single APL1 gene in *Anopheles stephensi*, as predicted by bioinformatic assembly and analysis short-read sequencing. Primers used for PCR sizing and for sequencing are indicated in Additional File 1: Figure S1. The annotated genes in the VectorBase genome database (24) corresponding to *A. stephensi* APL1 are ASTE016290 in *A. stephensi* SDA-500 strain, and ASTEI02571 in *A. stephensi* Indian strain. The Vectorbase assemblies and annotations used, current as of January 2019 were: SDA-500 strain, assembly AsteS1, gene set: AsteS1.7, dated 22 Oct 2018; and Indian strain, assembly AsteI2, gene set AsteI2.3, dated 21 Feb 2017.

For phylogenetic analysis of APL1 copy number as presented in Additional File 2: Figure S2, APL1C orthologues were obtained from Vector Base and sequence was extracted for +/− 60,000 base pairs (bp) around the APL1C orthologue and its flanking sequence. Sequences were compared and visualized in a pair-wise fashion using the tBlastX algorithm within the Double Act interface of the Artemis Comparison Tool (25) and visualized using Easyfig (26) to illustrate the number of APL1 family genes across species. Forward and reverse matches have been colored the same and percent ID cutoffs have been set to a minimum of 50% (light pink in Additional File 2: Figure S2 represents a 50% match and bright red 100% match, areas with less than 50% match are not depicted in color). Each mosquito species was compared directly to the *A. gambiae* PEST genome, the most mature *Anopheles* genome and the genome for which the APL1 gene family was originally annotated (16).

### Mosquitoes

*Anopheles stephensi* SDA-500 strain was initiated in Pakistan (27) and *Anopheles coluzzii* Ngousso strain was initiated in Cameroon (28). Both strains are housed in the insectaries of the CEPIA platform at the Institut Pasteur, Paris. Mosquitoes were reared under standard conditions at 26°C and 80% relative humidity, with a 12 h light/dark cycle and continuous access to 10% sucrose solution in cotton pads (19).

*A. stephensi* samples used for APL1 population variation analysis were 6 individuals from a colony initiated at Chabahar, Iran in 2011, 6 individuals from a colony initiated at Bandar-Abbas, Iran in 2008 (both strains maintained at the Institut Pasteur of Iran), and 1 wild-caught individual from Bandar-Abbas. An ∼800 bp portion of the APL1 coding sequence was amplified from individuals using *A. stephensi* APL1 primers Ast40F, TCACCCTATCCCACAACGAT, and Ast06R, ACTGTCACCGTCACAACTGC. Amplicons of individuals were sequenced and variant calls were confirmed on both strands by visual examination of ABI sequence chromatogram trace files. *A. coluzzii* APL1 sequences were previously published, generated from the Ngousso colony (29) or wild population (30), and deposited in public archives.

### Gene silencing

Double-stranded RNA (dsRNA) specific for target genes was synthesized using the T7 Megascript kit (Ambion) as described (18) using indicated primers (Supplementary Table 1, Additional File 3). For each targeted gene, 500 ng of dsRNA (but not more than 207nl volume, depending on concentration) was injected into the thorax of cold-anesthetized 1 d-old female mosquitoes using a Nanoject II Auto-Nanoliter Injector (Drummond Scientific). Mosquitoes were injected with dsRNA specific for the target gene, or with control dsRNA containing the irrelevant sequence of GFP. The efficiency of gene silencing was monitored 4 d after dsRNA injection in pools of 8 mosquitoes as follows. After total RNA extraction, cDNA synthesis was performed using M-MLV reverse transcriptase and random hexamers (Invitrogen). For each sample, 1µg of total RNA was used in each of three independent cDNA synthesis reactions. Triplicates were pooled and used as template for qPCR analysis. Real-time PCR was performed using an ABI Prism 7900HT sequence detector (Applied Biosystems). Reactions were prepared in 20 μl volumes using SYBR Green PCR master mix (Applied Biosystems) and 900nM primers with three serial dilutions of cDNA, each dilution assayed in triplicate. Primers used for verification of gene silencing are indicated (Supplementary Table 1, Additional File 3). PCR conditions were 95°C for 10 min followed by 40 cycles of 95°C for 15 s, 55°C for 15 s and 60°C for 45 s. mRNA level was normalized to self (*A. stephensi* or *A. coluzzii*) ribosomal protein rpS7 mRNA in each sample, and each gene silencing condition was compared to the control treated with GFP dsRNA.

### *Plasmodium* infection and phenotyping

Mosquitoes were fed on mice infected with *Plasmodium yoelii* strain delta-p230p-GFP (31) at 8-12% parasitemia with mature gametocytes. For parasite development, mosquitoes were maintained at 24°C and 70% relative humidity on 10% sucrose or 10% sucrose supplemented with penicillin 62.5 µg/mL, streptomycin 100 µg/mL, and gentamicin 50 µg/mL. To measure *P. yoelii* infection, mosquito midguts were dissected at 8 d post-infection and oocysts were counted by fluorescence microscopy. Infection phenotypes measured were oocyst infection prevalence, which is the proportion of mosquitoes carrying ≥1 oocyst among the total number of dissected mosquitos, and oocyst intensity, which is the oocyst count in mosquitoes with ≥1 oocyst. Mosquito infection phenotypes were determined for at least two independent biological replicates of ≥30 dissected mosquitoes per replicate.

Differences in infection prevalence were statistically tested using the Chi-Square test, and analysis of oocyst intensity differences used the Wilcoxon signed rank non-parametric test. Statistical differences in prevalence and intensity were first tested independently for each replicate as described above and p-values were empirically determined using 100,000 Monte-Carlo permutations. Following independent statistical tests for each replicate, and when the direction of change of each independent replicate was concordant, the p-values from independent tests of significance were statistically combined using the meta-analytical approach of Fisher (32). All statistical analyses were done using R (33).

### Mosquito mortality curves

Mosquito mortality was monitored in cages of at least 50 mosquitoes, recorded every 2 days until all mosquitoes died. Treatment with dsRNA was done in 3 d old mosquitoes and recording of mortality began 4 d after dsRNA injection in 7 d old mosquitoes. Blood feeding with or without *P. yoelii* was done at 4 d after dsRNA injection in 7 d old mosquitoes, and recording of mortality began 3 d following the normal or infected bloodmeal in 10 d old mosquitoes. Beginning at adult emergence, mosquitoes were maintained with 10% sucrose, and in the case of antibiotic treatment, supplemented with penicillin 62.5 µg/mL, streptomycin 100 µg/mL, and gentamicin 50 µg/mL. Two to three replicates were performed for each condition tested. A Cox proportional hazards regression model was fitted to the data using treatments as predictor terms (34, 35).

## RESULTS

### Phylogeny of APL1 gene expansion from a unique ancestor

Recent *in silico* annotation of the *Anopheles stephensi* reference genome detected a single APL1 gene (21). This is in contrast to the Gambiae species complex, where APL1 is comprised of a family of three paralogs, APL1A, APL1B, and APL1C, with distinct roles in immunity (16, 18). Because assembly of short-read sequences can be problematic for paralogous families, we first wished to confirm the *in silico* single-gene model for *A. stephensi* APL1. We manually sized and sequenced the ∼9 kb APL1 locus, which closed sequence assembly gaps and verified the presence of a single APL1 gene in *A. stephensi* (Additional File 1: Figure S1).

We then examined the phylogeny of APL1 in 19 reference genomes from 18 *Anopheles* species, which includes two independent assemblies for *A. stephensi* (21, 36). A single APL1 gene was identified in 12 species, including *A. stephensi*, while the genome assemblies that include the Gambiae complex and *A. christyi* display an expanded APL1 gene family (Figure 1, Additional File 2: Figure S2). The Gambiae complex members each carry three APL1 paralogs, with the same locus structure as previously described for the sister taxa *A. gambiae* and *A. coluzzii* (16, 30). The African species *A. christyi*, the closest sequenced relative outside the Gambiae complex, contains at least two APL1 genes and likely a third, but resolution is limited because the *A. christyi* genome assembly is suboptimal, with an APL1 locus comprised of three unjoined contigs with intervening sequence gaps (Additional File 2: Figure S2).

**Figure 1.**
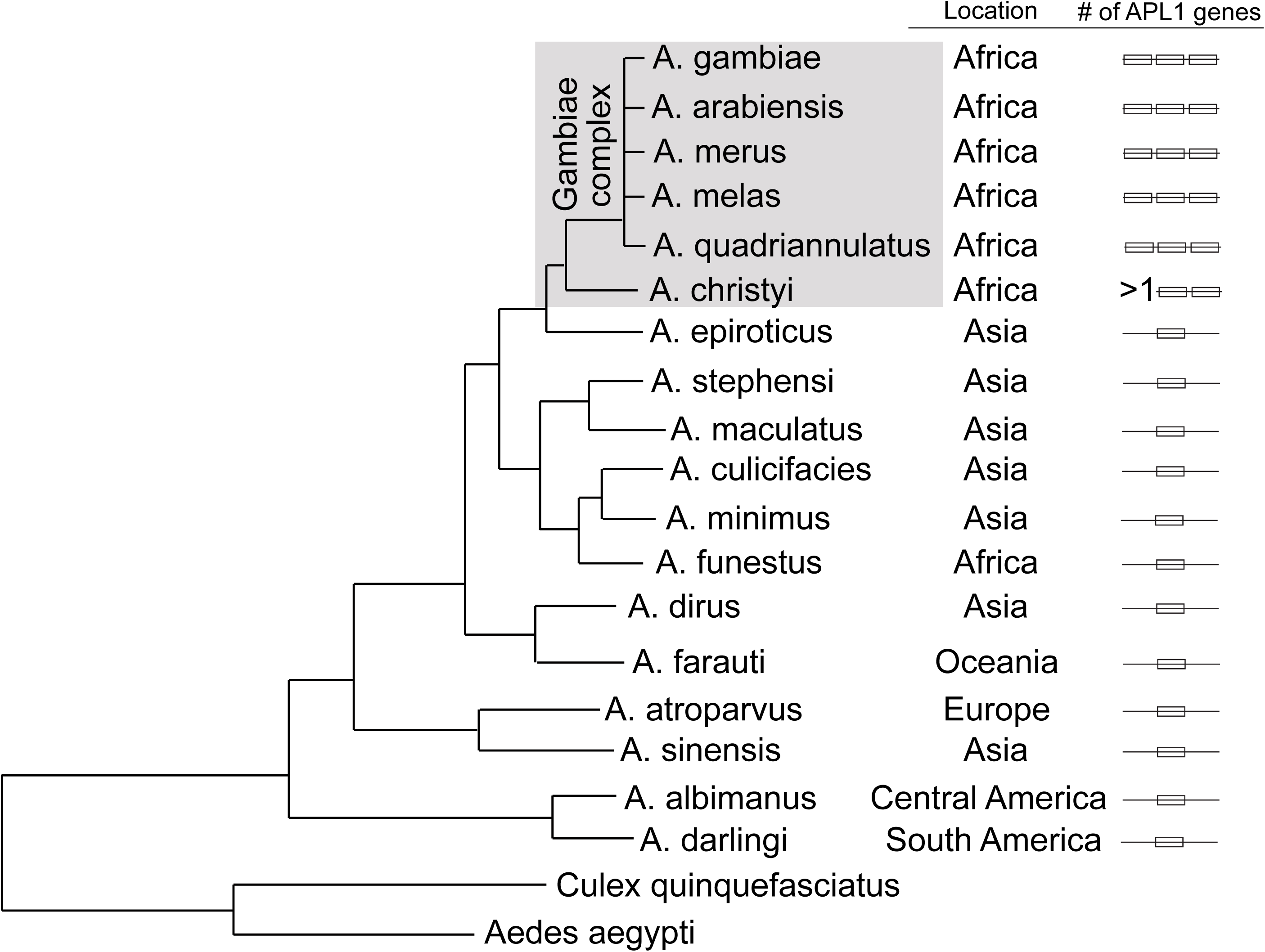
The APL1 gene underwent an expansion in an African *Anopheles* lineage. *Anopheles* phylogenetic tree indicates the number of APL1 gene paralogs present in the genome of 18 *Anopheles* species. Geographic locations of species and the number of APL1 genes in each species are indicated in columns, “Location” and “# of APL1 genes”, respectively. *Anopheles* species worldwide, including *A. funestus* in Africa, carry a single APL1 gene, which is the ancestral state. An exclusively African lineage displays an increased number of APL1 paralogs, including the Gambiae species complex and *A. christyi* (expanded APL1 lineage indicated by shaded box). The five sequenced Gambiae complex species clearly carry three APL1 paralogs, while *A. christyi* carries more than one and possibly three, but the genome assembly is poor, thus indicated as >1 APL1 gene. Phylogeny modified from (36).

The next most related sequenced relative to *A. christyi*, the Asian species *A. epiroticus*, carries a single APL1 gene (Figure 1, Additional File 2: Figure S2). Based on synteny, and the presence in *A. epiroticus* of a homolog of the gene AGAP007034 (located between *A. gambiae* APL1B and APL1C), the single APL1 gene in *A. epiroticus* displays the greatest relatedness to *A. gambiae* APL1C, with APL1B and APL1A presumably arising through duplication events during divergence of the Gambiae complex and *A. christyi* from their common ancestor. The *Anopheles* species carrying an expanded APL1 gene complement do not correspond precisely to the monophyletic Pyretophorus taxonomic group of *Anopheles* species (37, 38). The Pyretophorus group includes *A. christyi* and the Gambiae complex, which carry an expanded APL1 locus, and also *A. epiroticus*, which has only one APL1 gene. Outside the group of *A. christyi* and the Gambiae complex, the evidence clearly supports a unique APL1 gene in all species, with the possible exception of *A. minimus*, which is a poor quality assembly and consequently the possibility of two APL1 genes cannot be excluded (Additional File 2: Figure S2). Thus, we conclude that the single APL1 gene found in most sequenced *Anopheles* including *A. stephensi* represents the ancestral state of this locus, while the expansion of APL1 to three genes is a derived state, restricted to the Gambiae complex and *A. christyi*.

### APL1 population variation

Genetic polymorphism within the unique APL1 gene in *A. stephensi* was measured by sequencing of individual mosquitoes colonized from the natural population in Iran. The unique ancestral APL1 gene in these mosquitoes segregates 7 SNP sites over 1190 bp, or ∼6 variable nucleotide sites per kilobase (kb). By comparison, the APL1C paralog in *A. coluzzii* mosquitoes (the Ngousso colony from Cameroun) measured in the same way segregates 117 SNP sites in 2924 bp, or ∼40 variable sites per kb (29), which is more than six-fold greater polymorphism than the unique *A. stephensi* APL1 gene. *A. stephensi* APL1 is compared to *A. coluzzii* APL1C because APL1C displays the closest orthology to the unique APL1 (Additional File 2: Figure S2). However, in the natural West African population of *A. gambiae* and *A. coluzzii*, paralog APL1A is even more polymorphic than APL1C, displaying approximately double the diversity (30). The differing levels of diversity of the unique APL1 ancestor and the three APL1 paralogs suggest the genes are exposed to distinct natural selection, likely due to functional differences, and greater evolutionary constraint in the case of the single ancestral APL1 gene.

### Depletion of *A. stephensi* APL1 reduces mosquito lifespan

Depletion of APL1 in *A. stephensi* by RNAi-mediated silencing led to significantly elevated mosquito mortality as compared to controls treated with an irrelevant double-stranded RNA (dsRNA), dsGFP. The effect was seen regardless of whether APL1 depletion was followed by a sugar meal or bloodmeal (Figure 2A and 2B), and the reduction of mosquito lifespan was even more pronounced when APL1 silencing was followed by a *Plasmodium yoelii*-infective bloodmeal (Figure 2C). After parasite infection, ∼70% of APL1-depleted mosquitoes died by 8 d post-infection as compared to ∼15% mortality in the dsGFP-treated controls.

**Figure 2.**
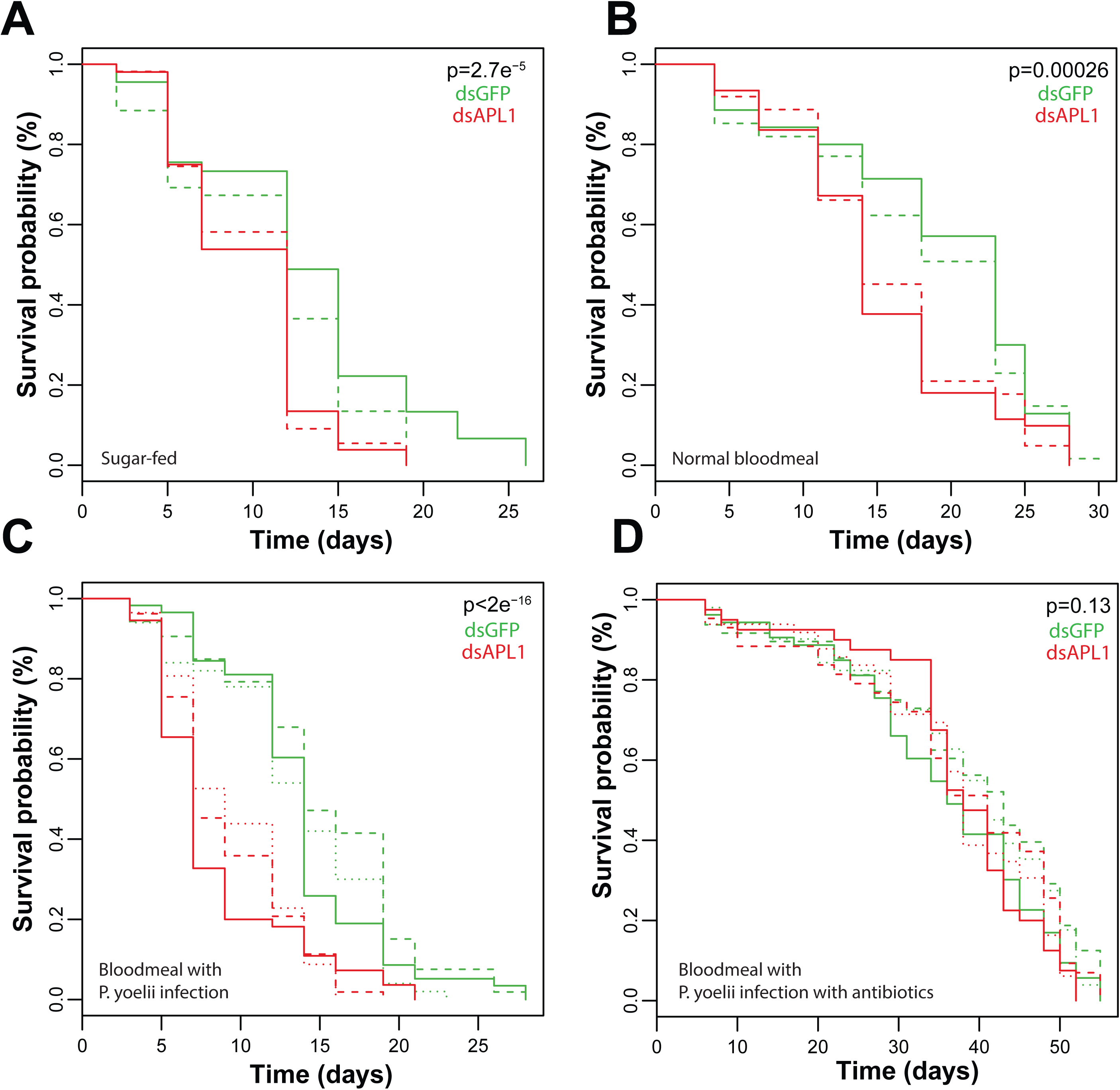
Depletion of APL1 leads to mosquito mortality in *Anopheles stephensi*. Survival curves of *A. stephensi* depleted for APL1 activity by dsAPL1 treatment (red lines) as compared to dsGFP-treated controls (green lines) under different experimental conditions**. A)** Sugar-fed mosquitoes. **B)** Mosquitoes fed an uninfected normal bloodmeal. **C)** Mosquitoes fed a *Plasmodium yoelii*-infected bloodmeal. **D)** Mosquitoes treated with antibiotics and fed a *P. yoelii*-infected bloodmeal. Replicate experiments are distinguished by line type (plain, dashed or dotted, respectively). X-axis indicates time after the start of recording. A Cox proportional hazards regression model was fitted to the data using treatment and replicate as predictor terms. The p-value associated with the dsRNA treatment term of the Cox model is shown on each panel.

### Elevated mortality of APL1-depleted *A. stephensi* is rescued by antibiotic treatment

The observed mortality after depletion of an immune gene suggested a potential antibacterial activity for APL1 in *A. stephensi*. The additional mortality after parasite infection would be consistent, because malaria ookinete invasion from the midgut lumen facilitates physical entry of bacteria into epithelial cells and heightens microbial exposure (39). This observation suggests that the bacteria leading to elevated mortality in APL1-depleted *A. stephensi* are likely of enteric origin. The three APL1 paralogs found in the Gambiae complex are known to mediate protection from *Plasmodium* infection (19), but their involvement in protection against other pathogens including bacteria has not been reported.

To test the hypothesis that *A. stephensi* APL1 protects from potentially pathogenic bacteria, newly emerged adult *A. stephensi* mosquitoes were fed antibiotics in the sugar meal, were then treated with dsAPL1 or dsGFP, and were infected with *P. yoelii* parasites. Antibiotic feeding abolished the elevated mortality associated with loss of APL1 function, even in the most pronounced case of *Plasmodium* infection (Figure 2D). These results suggest that the activity of APL1 is essential to protect from bacteria under a range of biological conditions in *A. stephensi*, and that loss of APL1 function leaves the host vulnerable to lethal bacterial effects.

### Simultaneous depletion of all three APL1 paralogs does not reduce *A. coluzzii* lifespan

In contrast to the elevated mortality observed in APL1-depleted *A. stephensi*, a mortality effect has not been reported for the APL1 paralogs in *A. gambiae* and *A. coluzzii* (13, 16, 18–20). To confirm this apparent phenotypic difference between the ancestral and expanded APL1 genes, we tested the effect of loss of all APL1 activity in *A. coluzzii* by depleting all three APL1 paralogs. Simultaneous depletion of all three APL1 paralogs did not alter longevity of *A. coluzzii* after sugar feeding (Figure 3A) or *Plasmodium* infection (Figure 3B). Thus, different from the single APL1 gene in *A. stephensi*, which displayed elevated mortality under these conditions, activity of the three APL1 paralogs in *A. coluzzii* are not required for protection against bacteria.

**Figure 3.**
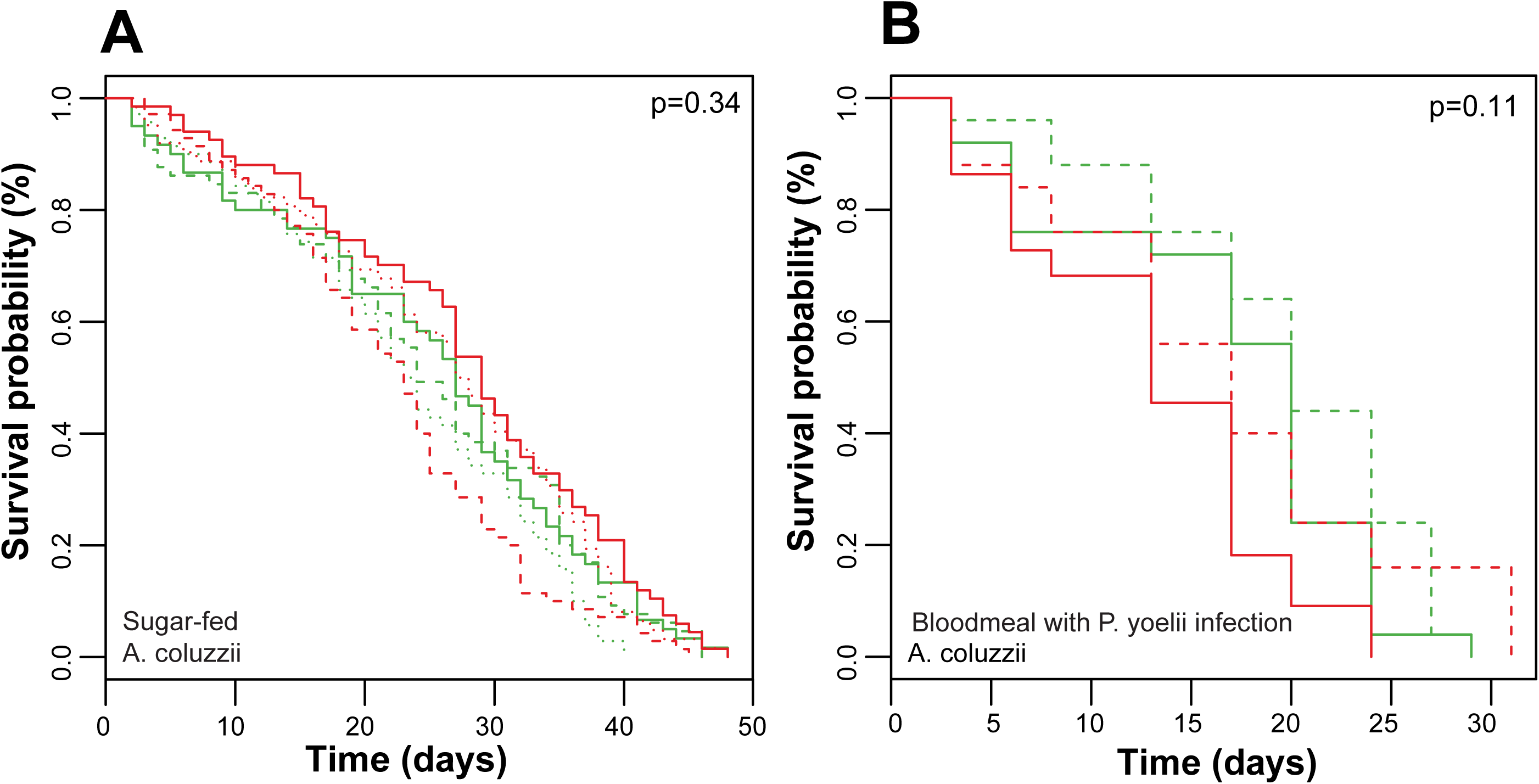
Simultaneous depletion of all three APL1 paralogs in *A. coluzzii* does not cause mosquito morality. **A)** Survival curves of *A. coluzzii* depleted for APL1 activity by dsAPL1 treatment (red lines) as compared to dsGFP controls (green lines), for sugar-fed mosquitoes and **B)** survival curves for mosquitoes fed a *Plasmodium yoelii*-infected bloodmeal. Survival curves from replicates are distinguished by line type (plain, dashed or dotted, respectively). X-axis indicates time after the start of recording, not mosquito age (see Methods). A Cox proportional hazards regression model was fitted to the data using treatment and replicate as predictor terms. The p-value associated with the dsRNA treatment term of the Cox model is shown on each panel.

### Anti-*Plasmodium* protection by *A. stephensi* APL1 is secondary to its antibacterial function

Depletion of the unique APL1 gene in *A. stephensi* led to decreased *P. yoelii* parasite load (Figure 4A). However, the APL1-depleted *A. stephensi* were already compromised due to elevated bacteria-derived effects and mortality, and may have been physiologically unable to support *Plasmodium* development. When bacteria are controlled by antibiotics, though, APL1-depleted *A. stephensi* carried significantly greater *P. yoelii* infection loads as compared to dsGFP-treated controls (Figure 4B). Thus, controlling for the antibacterial effect of APL1 in *A. stephensi* revealed an underlying anti-*Plasmodium* activity of the unique APL1 gene, but the dominant function of the gene appears to be control of bacteria that are often lethal without protection.

**Figure 4.**
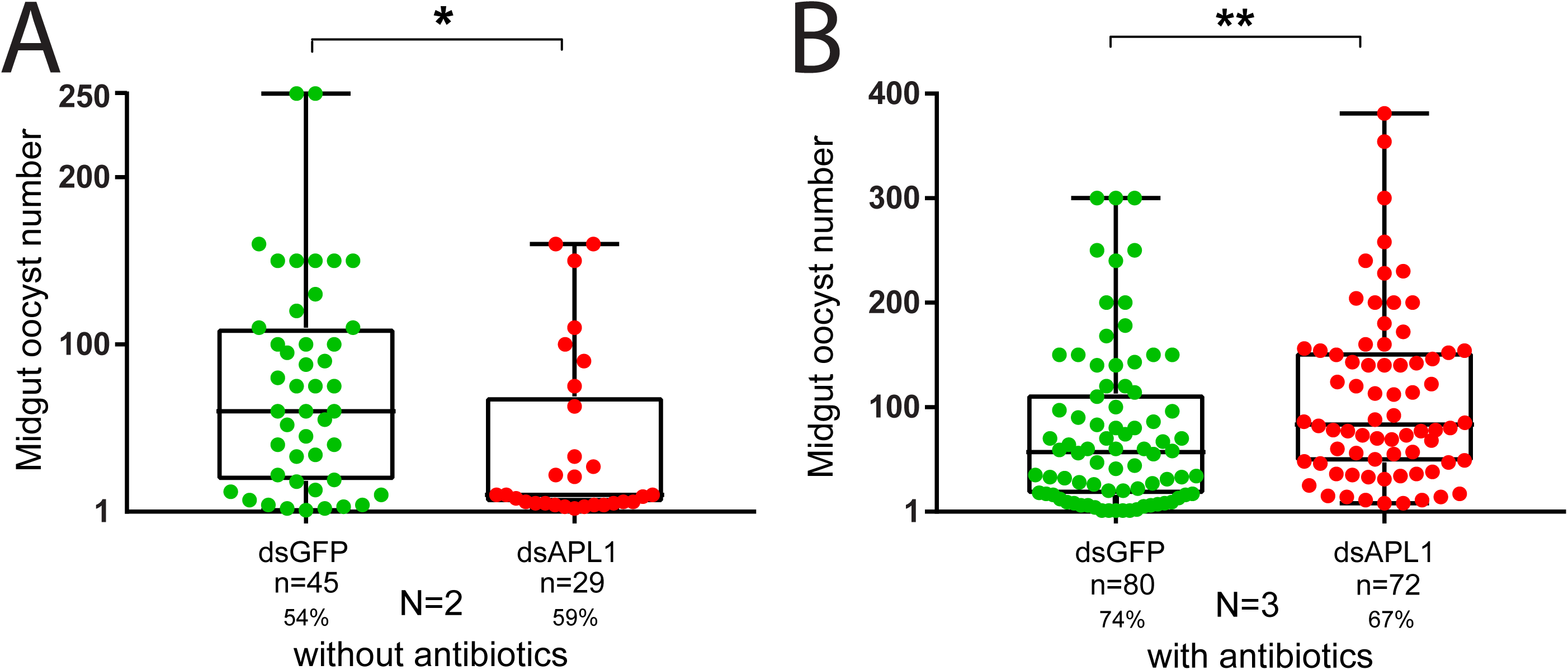
*Anopheles stephensi* APL1 protection from *Plasmodium yoelii* infection is secondary to its antibacterial function. **A)** *P. yoelii* oocyst infection intensity in *A. stephensi* mosquitoes treated with dsAPL1 or control dsGFP, both without antibiotic treatment. Oocyst intensity is the oocyst count in mosquitoes with ≥1 oocyst, to avoid confounding with infection prevalence. Oocyst infection prevalence, the proportion of mosquitoes carrying ≥1 oocyst, is indicated as a percentage below sample sizes. Number of biological replicates is indicated below plots. **B)** As in A, but mosquitoes were subject to antibiotic treatment before *Plasmodium* exposure. Significance levels of combined p-values: * p-value<0.05; ** p-value <0.01.

These results are in contrast to silencing of the three APL1 paralogs In *A. coluzzii*, which consistently leads to elevated *Plasmodium* infection levels (18, 19) but not elevated mortality (Figure 3). Therefore, the three APL1 paralogs confer protection against *Plasmodium* infection independently of the need to protect against bacteria. Taken together, these results indicate that the combined phenotype of the three paralogs does not recapitulate the phenotype of the ancestral single gene. This suggests that the divergence of the three APL1 paralogs from the unique APL1 ancestor was accompanied by important functional changes. Protection from bacteria in the expanded-APL1 mosquito lineage may have been functionally replaced by other unknown immune factors or distinct physiological mechanisms that protect from bacteria or their pathogenic effects.

### The unique APL1 gene in *A. stephensi* displays an ancestral immune signaling profile

The APL1 paralogs in *A. coluzzii* are transcriptionally regulated by distinct immune signaling pathways. Expression of paralog APL1A is regulated by the transcription factor Rel2, the positive regulator of the Immune deficiency (Imd) immune pathway, while paralog APL1C is regulated by transcription factor Rel1, positive regulator of the Toll pathway (16, 18, 19, 40).

We tested the effect of these signaling pathways on expression of the unique APL1 gene in *A. stephensi*. Activation of Toll signaling in *A. stephensi* by depletion of the Toll negative regulator, Cactus (Figure 5A), led to increased APL1 expression (Figure 5B), and depletion of the Imd positive regulator Rel2 (Figure 5C) led to reduced APL1 expression in *A. stephensi* (Figure 5D). Consequently, *A. stephensi* APL1 expression is under control of both the Toll and Imd pathways. A previous study found that overexpression of a Rel2 transgene in *A. stephensi* induced APL1 expression, consistent with our findings, but the response to Rel1 was not tested (22). Thus, the ancestral unique APL1 gene in *A. stephensi* is regulated by two signaling pathways, Toll and Imd, while after APL1 gene duplication and divergence, these two controlling pathways were subdivided to the derived paralogs APL1C and APL1A, respectively.

**Figure 5.**
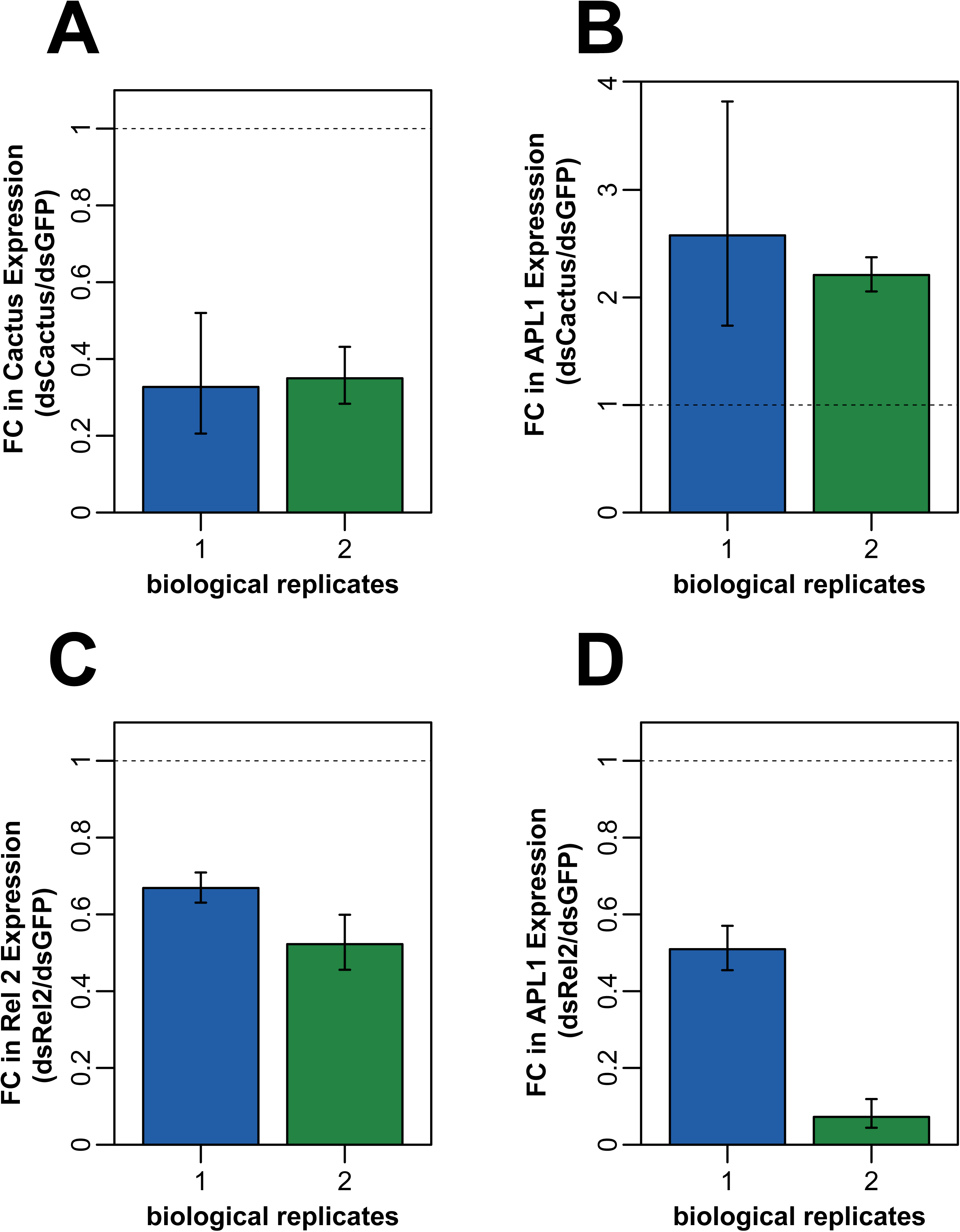
Transcription of *Anopheles stephensi* APL1 is regulated by both Toll and Imd immune signaling pathways. Regulation of expression of the unique *A. stephensi* APL1 gene was queried by silencing the negative regulator of Toll, Cactus (A and B) or the positive regulator of Imd, Rel2 (C and D). **A)** Cactus expression is efficiently suppressed by treatment with double-stranded RNA targeting Cactus (dsCactus). **B)** APL1 expression is augmented by silencing of Cactus, which constitutively activates the Toll pathway. Fold change of APL1 gene expression in *A. stephensi* depleted for Cactus by dsCactus treatment, relative to dsGFP treated controls. **C)** Rel2 expression is efficiently suppressed by treatment with dsRNA targeting Rel2 (dsRel2). **D)** APL1 expression is diminished by silencing of Rel2, which inhibits Imd pathway activity. Fold change of APL1 gene expression in *A. stephensi* depleted for Rel2 by dsRel2 treatment, relative to dsGFP treated controls. Transcript abundance measured by qPCR in two biological replicates as indicated.

## DISCUSSION

We find that *A. stephensi* and most other sequenced *Anopheles* species carry a unique APL1 gene, which expanded to three paralogs in an exclusively African lineage that includes (of sequenced species) just *A. christyi* and all members of the Gambiae complex. Activity of the unique APL1 gene is required in *A. stephensi* for protection from bacteria. The function of the unique gene is distinct from the expanded APL1 gene family, which protect against at least *Plasmodium* but are not essential for protection from bacteria. The unique APL1 gene displays the ancestral immune signaling profile, because its expression is regulated by both Toll and Imd pathways, in distinction to the paralogs in *A. coluzzii*, in which regulation by the immune pathways evolved to be specialized to different gene family members.

### Function of ancestral and derived APL1 genes

The unique APL1 gene is essential for *A. stephensi* fitness and survival, while the three paralogs combined are not essential for *A. coluzzii* under the same conditions, because their depletion does not have lethal consequences. Gene essentiality is dependent on genomic and biological context, including environmental conditions (41). The common ancestor of the Gambiae complex-*A. christyi* lineage evolved to exploit an unknown ecological niche, probably in Africa because all species known to carry an expanded APL1 locus are African, and may have encountered new environmental pathogens (30, 42, 43). The simplest interpretation is that essentiality of the ancestral unique APL1 gene was lost at the time of the expansion and functional divergence of the three paralogs. The expanded APL1 paralogs evolved new immune roles, exemplified by observed functional differences among the three paralogs in the Gambiae complex (18–20). However, the paralogs did not simply subdivide the functions of the unique ancestor, because they are not required for protection against bacteria under the conditions tested. The expansion of the APL1 gene family was likely accompanied by a suite of other unknown genomic changes necessary for adaptation of the Gambiae complex-*A. christyi* lineage to the new ecological niche. Protection from bacteria was presumably shifted to other unknown genes or physiological factors, which may have evolved at the same time.

Previous population sequencing revealed that the three APL1 paralogs in the Gambiae complex are exceptionally polymorphic and display signals of adaptive maintenance of variation, including maintenance of alleles that are older than the species within the Gambiae complex (30). This genetic pattern is consistent with a model of balancing polymorphism maintained by exposure to fluctuating environmental pathogens in a trench warfare dynamic (44). In contrast, examination of *A. stephensi* APL1 sequences from individual mosquitoes from the Iran population suggests that diversity of the unique APL1 gene is quite low. The simplest model is that unique APL1 is under selection mainly to protect the host from relatively stable groups of enteric bacteria. Additional population resequencing will be required to test these hypotheses.

### APL1 copy number and malaria vectorial capacity

Expanded APL1 copy number does not directly correlate with DVS status, but this comparison is confounded with mosquito behavior, because not all expanded-APL1 species are human feeding. The four expanded-APL1 species that are DVS display high human-biting preference (*A. gambiae*, *A. coluzzii*, *A. merus*, *A. melas*), while the other two sequenced species with an expanded APL1 locus, the non-vectors *A. christyi* and *A. quadriannulatus*, are cattle-feeding species in nature (6, 45). Of these latter two non-vector species, *A. quadriannulatus* is physiologically susceptible to infection with *P. falciparum* when fed experimentally on parasitemic blood (6, 7), and vector competence of *A. christyi* has not been tested.

The more interesting question is whether, among human-feeding species, carriage of the expanded APL1 locus influences the efficiency of malaria transmission. The human-feeding members of the Gambiae complex are considered the most efficient malaria vectors in the world (46, 47), and all of these species carry the expanded APL1 locus. Their efficient malaria transmission could be a secondary consequence of inhabiting African ecological niches that also happen to be particularly favorable to malaria transmission (9, 10, 12). However, other African vectors such as *A. funestus*, *A. nili*, *A. pharoensis* and *A. moucheti* are DVS, but are often described as locally important secondary vectors, and lack the epidemiological impact of the expanded-APL1 Gambiae complex. *A. funestus* carries a single APL1 gene, and *A. nili*, *A. pharoensis* and *A. moucheti* have not been sequenced but based on the phylogenetic analysis are also expected to carry the single ancestral APL1 locus.

Thus, the current results raise the question whether the correlation of APL1 copy number with vectorial efficiency is accidental or biologically meaningful. The ancestral single APL1 protects *A. stephensi* against malaria parasite infection, but this activity is secondary to a dominant and essential function of protection against bacteria. Under these conditions, it would not seem adaptive for *Plasmodium* to inhibit the activity of the unique APL1 in order to modulate anti-malaria immunity, because parasite transmission requires vector survival for ≥14 days. Thus, parasite inhibition of unique APL1 immune function in *A. stephensi* would be expected to decrease vector survival and therefore the parasite’s own reproductive fitness. In contrast, in *A. coluzzii* with three APL1 paralogs, malaria immunity and protection from bacteria are uncoupled, because loss of APL1 function does not reduce longevity. The separation of anti-*Plasmodium* immunity from antibacterial protection should allow *Plasmodium* (and other pathogens) to subvert APL1-mediated immunity without the risk of provoking host mortality.

## Conclusions

The ancestral and derived APL1 loci, represented by *A. stephensi* and *A. coluzzii*, respectively, display large differences in essentiality, function, gene regulation and genetic diversity. Manipulative experimentation and population genetic analysis will be required to understand the functional and ecological significance of the ancestral and derived APL1 for immunity and malaria transmission.

## Abbreviations

bp: base pairs
d: days
dsRNA: double strand RNA
DVS: dominant malaria vector species
RNAi: RNA interference

## DECLARATIONS

### Ethics approval and consent to participate

The protocol for the ethical treatment of the animals used in this study was approved by the research animal ethics committee of the Institut Pasteur, “C2EA-89 CETEA Institut Pasteur” as protocol number B75-15-31. The Institut Pasteur ethics committee is authorized by the French Ministry of Higher Education and Research (MESR) under French law N° 2001-486, which is aligned with Directive 2010/63/EU of the European Commission on the protection of animals used for scientific purposes.

### Consent for publication

Not applicable.

### Availability of data and materials

All sequence files are available from the EBI European Nucleotide Archive database (http://www.ebi.ac.uk/ena/) under ENA study accession number [REQUESTED].

### Author contributions

Designed research: CM, EB, KE, NDD, NS, MMR, KDV

Performed research: CM, EB, KE, IH, CD, EBF, AR, SZ, MIKN, NDD, MMR

Analyzed data: CM, EB, KE, IH, CD, AR, SZ, MIKN, MMR, KDV

Wrote the paper: CM, EB, KE, IH, NDD, MMR, KDV

All authors read and approved the final manuscript.

## Acknowledgments

We thank the Institut Pasteur core facility, the Center for the Production and Infection of *Anopheles* (CEPIA), for gametocyte culture and mosquito infection, and Corinne Genève, GGIV Institut Pasteur, for rearing and infecting mosquitoes. We thank Olivier Sylvie, Inserm unit on Molecular biology and immunology of malaria liver infection, for provision of *P. yoelii* fluorescent strain delta-p230p-GFP.

## SUPPORTING FILE CAPTIONS

**Additional File 1: Figure S1.**
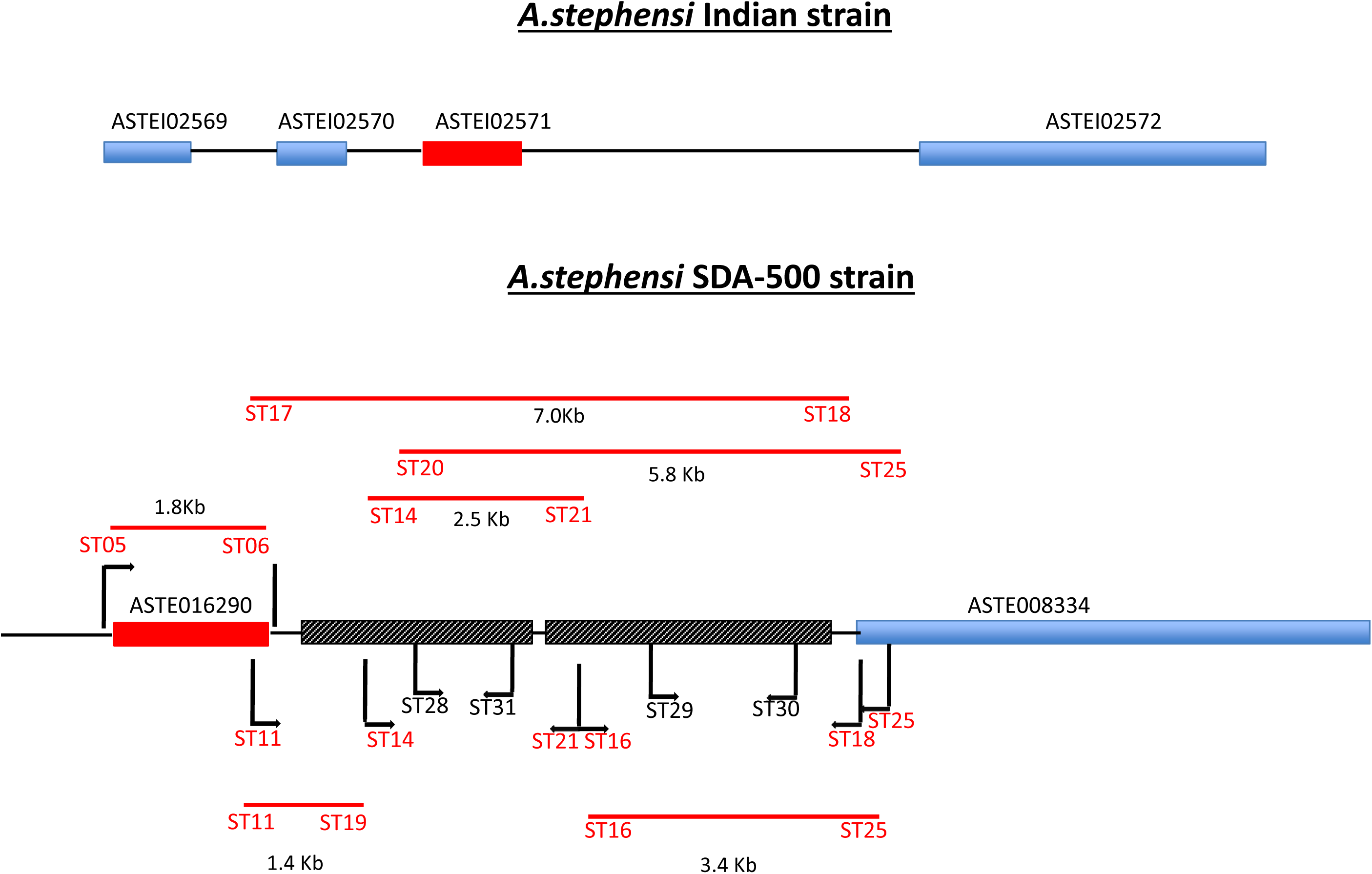
Manual sequence confirmation of unique APL1 gene in *Anopheles stephensi* strain SDA-500. PCR sizing reactions and manual sequencing of the *A. stephensi* SDA-500 strain APL1 locus. Primer locations used for sizing, generation of amplicon templates for sequencing, and for sequence walking along amplicon templates for are indicated with code “ST” and number, primer sequences given in Supplementary Table 1, Additional File 3. The *A. stephensi* APL1 ortholog is indicated as a red rectangle (labelled as ASTE016290 in strain SDA-500 and ASTEI02571 in strain Indian). Unjoined sequence gaps in the SDA-500 strain assembly are indicated by hatched rectangles. Manual sizing was done by PCR, and resulting fragment sizes generated by indicated primer pairs are presented as horizontal lines over the relevant interval. The large amplicons were manually Sanger-sequenced as uncloned amplicons using the indicated primers for walking along the amplicon. The integration of sizing and sequencing resolved the unjoined region in the assembled SDA-500 sequence, and eliminated the possibility that other APL1 paralogs are concealed in the hatched rectangles.

**Additional File 2: Figure S2.**
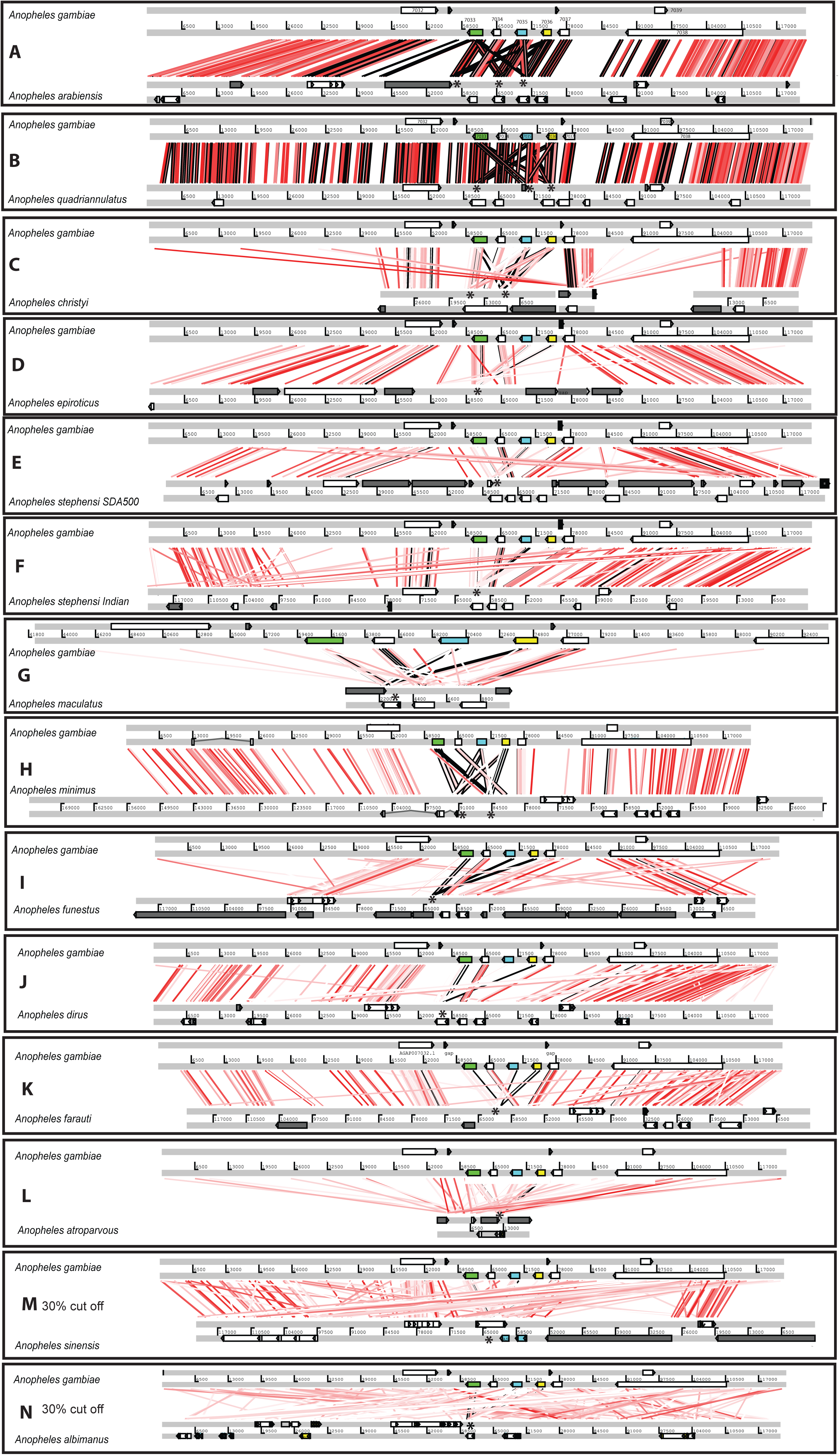
APL1 gene copy number in *Anopheles* genome sequence assemblies. The orthologs of APL1C, the most basal of the three expanded APL1 paralogs, were obtained from mosquito genome assemblies in VectorBase (24). Sequences were compared to *A. gambiae* in a pair-wise fashion to determine the number of APL1 family genes by phylogenetic comparison using the Double Act interface of the Artemis Comparison Tool (25) and the tBlastX algorithm, visualized using Easyfig (26). The *A. gambiae* reference is shown at the top of each pairwise comparison panel. Forward and reverse matches are colored the same and percent ID cutoffs were set to a minimum of 50% (light pink represents a 50% match and bright red 100% match, areas with less than 50% match are not depicted in color), except in the bottom two panels (N and O) a 30% ID cutoff was used. **A) *A. arabiensis***, an African Gambiae complex malaria vector species. **B) *A. quadriannulatus***, an African Gambiae complex non-vector species. **C) *A. christyi***, an African non-vector related to the Gambiae complex (boundaries of three unconnected scaffolds of the *A. christyi* assembly in this region are indicated by white space). **D) *A. epiroticus***, an Asian malaria vector. **E) *A. stephensi* SDA-500 strain**, an Asian malaria vector. **F) *A. stephensi* Indian strain**, an Asian malaria vector. **G) *A. maculatus***, an Asian malaria vector. **H) *A. minimus***, an Asian malaria vector. **I) *A. funestus***, an African malaria vector. **J) *A. dirus***, an Asian malaria vector, **K) *A. farauti***, an Oceania malaria vector, **L) *A. atroparvus***, a European malaria vector, **M) *A. sinensis***, an Asian malaria vector and **N) *A. albimanus***, a Central American malaria vector. Predicted genes depicted as white rectangles, the APL1 gene family in *A. gambiae* are colored rectangles: APL1C (AGAP007033), green; APL1B (AGAP007035), blue; APL1A (AGAP007036), yellow. Unjoined sequence gaps of one or more N positions, grey rectangles.

**Additional File 3: Table S1. Sequence of primers**. Name preceded by “ST”, primers used for manual sequence annotation of APL1 locus (Additional File 1: Figure S1). Sequence of the primers (5’-3’) used for the synthesis of double-stranded RNA for gene silencing (name preceded by “T7”), and for verification of silencing efficiency by quantitative RT-PCR (name preceded by “Ast”).

